# Effects of anthropogenic wildfire in low-elevation Pacific island vegetation communities in French Polynesia

**DOI:** 10.1101/196352

**Authors:** Erica A. Newman, Carlea A. Winkler, David H. Hembry

**Author notes:** Corresponding author: Erica A. Newman.

## Abstract

Anthropogenic (or human-caused) wildfire is an increasingly important driver of ecological change on Pacific islands including southeastern Polynesia, but fire ecology studies are almost completely absent for this region. Where observations do exist, they mostly represent descriptions of fire effects on plant communities before the introduction of invasive species in the modern era. Understanding the effects of wildfire in southeastern Polynesian island vegetation communities can elucidate which species may become problematic invasives with continued wildfire activity. We investigate the effects of wildfire on vegetation in three low-elevation sites (45-379 m) on the island of Mo’orea in the Society Islands, French Polynesia, which are already heavily impacted by past human land use and invasive exotic plants, but retain some native flora. In six study areas (3 burned and 3 unburned comparisons), we placed 30 transects across sites and collected species and abundance information at 390 points. We analyzed each local community of plants in three categories: natives, those introduced by Polynesians before European contact (1767 C.E.), and those introduced since European contact. Burned areas had the same or lower mean species richness than paired comparison sites. Although wildfire did not affect the proportions of native and introduced species, it may increase the abundance of introduced species on some sites. Non-metric multidimensional scaling indicates that (not recently modified) comparison plant communities are more distinct from one another than are those on burned sites. We discuss conservation concerns for particular native plants absent from burned sites, as well as invasive species (including *Lantana camara* and *Paraserianthes falcataria*) that may be promoted by fire in the Pacific.

## Introduction

The ecological effects of fire in tropical Pacific island ecosystems have received little attention in most Pacific island groups (Nunn 1990). Despite the potential importance of wildfires for conservation, public health, and human-environment interactions, very few studies of wildfire effects on Pacific island vegetation exist, and reliable, quantitative information on conservation threats is limited (Keppel *et al.* 2004). Almost nothing is known about how contemporary wildfire affects plant communities in the Pacific outside of Hawai‘i (Hughes *et al.* 1991; D’Antonio and Vitousek 1992; Hughes and Vitousek 1993; Freifelder *et al.* 1998; D’Antonio *et al.* 2000, 2001; Ainsworth and Kauffman 2009, 2010, 2013; Angelo and Daehler 2013; Ellsworth *et al.* 2014), with a few studies in New Caledonia (McCoy *et al.* 1999), western Polynesia (Hjerpe *et al.* 2001; Elmqvist *et al.* 2002; Franklin 2007) and Guam (Athens and Ward 2004). Furthermore, human-caused wildfires are increasingly occurring on Pacific islands in the modern era (Trauernicht *et al*. 2015), and islands are especially susceptible to anthropogenic disturbances (Keppel *et al.* 2014). In this paper, we use a novel dataset to examine the effects of wildfires on low-elevation and already heavily-invaded vegetation communities in the Society Islands, French Polynesia.

Few records and no formal studies exist for French Polynesia of the ecological effects of wildfire in modern times. Indeed, most Pacific island plant communities, especially communities dominated by native plant species, are not considered to support active fire regimes of the type seen in some continental communities of similar climate due to differing ignition patterns, plant structure, winds, and other factors (*e.g.* Cochrane 2003; but see Hunter-Anderson 2009). Although natural wildfire is thought to be extremely rare as a disturbance in Pacific terrestrial native plant communities, botanists working in the region have long suspected that anthropogenic fire (alongside other human land-use changes) has played an important role in certain low-and mid-elevation terrestrial vegetation communities. The botanical evidence for a role of anthropogenic fire are the extensive low-elevation fernlands and grasslands which drew the attention of early Western field botanists in the Pacific (including French Polynesia). These communities were usually described as near-monocultures, dominated by either the native fern *Dicranopteris linearis* (Gleicheniaceae) or several species of grass. In some contemporary cases, these fernlands and grasslands take the form of “savannas” with scattered emergent trees in genera such as *Metrosideros* and *Commersonia* (Florence 1997). For example, *Dicranopteris* fernlands are common in the Society, Cook, Austral, Hawaiian, and Wallis and Futuna islands (Florence 1997; Sykes 2016; Meyer 2017). In a detailed survey of Society Islands vegetation published in French, Papy (1954) reported an association of *Dicranopteris* and grasses with fire, and suggested that the former can resprout from rhizomes following wildfire. He also observed that certain woody plants (*Metrosideros collina, Melastoma denticulatum*, and *Dodonaea viscosa*) are often associated with areas that have burned. These observations are consistent with more recent ones from Hawai‘i for *Dodonaea viscosa* and the congeneric *Metrosideros polymorpha* (following single, low-intensity fires; Ainsworth and Kauffman 2009; D’Antonio *et al.* 2000). Surveying the vegetation of the entire tropical Pacific many decades later, Mueller-Dombois and Fosberg (1998) came to a similar conclusion, arguing that *Dicranopteris* fernlands represent potentially anthropogenic post-disturbance communities because of their extremely low plant diversity, and because no plant species other than *D. linearis* are restricted to these communities (*i.e.* these communities contain no habitat specialists).

Pacific archaeologists and paleoecologists have concurred with the conclusions of botanists, invoking charcoal and other pollen records as further evidence that fire is anthropogenic (Kahn *et al.* 2015; Stevenson *et al.* 2017), and that widespread “pyrophytic” grassland and fernland zones resulted from the use of fire by the first colonizing humans in French Polynesia and elsewhere in the Pacific (Dodson and Intoh 1999; Kirch 2000; Athens and Ward 2004; Mann *et al.* 2008; but see Hunter-Anderson 2009). It is important to stress that local knowledge that anthropogenic fire promotes *Dicranopteris* savanna (*e.g*., references and informants cited in Meyer 2017) is consistent with these archeological and botanical interpretations, and may have been the original impetus which led Western investigators to suspect a role for fire in Pacific vegetation ecology. These independent lines of evidence point to anthropogenic fire in combination with other human impacts partially transforming the mid-and low-elevation vegetation of oceanic Pacific islands from forest to pyrophytic fernlands and grasslands, commencing perhaps as soon as the first human colonization of these islands (1000 C.E. for French Polynesia) (Mueller-Dombois and Fosberg 1998; Kirch 2000; Kahn *et al.* 2015; Stevenson *et al.* 2017).

The conclusions of botanists, archaeologists, and paleoecologists that fire in the Remote Pacific is primarily of anthropogenic origin and has played an important role ecologically only in transforming some low-and mid-elevation plant communities into fern and grass-dominated savannas are of particular importance for conservation and human welfare in French Polynesia, because anecdotal evidence suggests that wildfire events may be frequent during the 2000’s and 2010’s, and possibly increasing in frequency relative to historical baselines. Although comprehensive records are not publically available, published news reports in local media (*La Dépêche de Tahiti*, published daily in Papeete, Tahiti and the “Polynésie 1ère” news website) indicate wildfires events on the islands of Tahiti in 2013 (two fires), 2015, and 2016, Mo’orea in 2008 and 2011, Huahine in 2016, Raiatea in 2016, and Bora Bora in 2013 (Society archipelago) and Nuku Hiva in 2012 and Ua Pou in 2013 (Marquesas archipelago). A non-peer-reviewed study from a field course taught by the University of California, Berkeley identified three wildfires on Moorea which occurred in 1991, 1995, and 1998 (Koehler 1999), and partial wildfire records held by the Sapeurs-Pompiers de Moorea (the Moorea fire department) list 5 wildfires occurring in the first half of 2011 (J.-M. Hokaupoko, *pers. comm*.). Despite the potential importance of wildfire as a problem for both conservation (J.-Y. Meyer, *pers. comm*.) and human well-being (J.-M. Hokaupoko, *pers. comm*.) in French Polynesia, no published study has examined the ecological effects of wildfire in that country.

The absence of studies examining ecological consequences of wildfire in French Polynesia are concerning because analogous studies from the Hawaiian Islands (which are ecologically similar and share many native plant genera) suggest that wildfires may actually have very serious effects on modern vegetation communities in ways which impact both biodiversity and the propensity for future wildfires. Studies of wildfire on the island of Hawai‘i (“the Big Island”) have shown that although a considerable number of native woody and tree fern species in wet and mesic forests show some ability to persist after wildfire through resprouting or germination following lava-ignited wildfire (Ainsworth and Kauffman 2009), and some native herbaceous species appear to sprout only following repeated wildfires that destroy litter (Ainsworth and Kauffman 2013), most evidence suggests that wildfires increase abundance of introduced herbaceous and grass species. Wildfires promote invasion of introduced herbaceous species in forests (Ainsworth and Kauffman 2010), native tree and tree fern death (D’Antonio *et al.* 2000; Ainsworth and Kauffman 2013), and the replacement of native communities with invasive grasslands (Hughes *et al.* 1991; Hughes and Vitousek 1993; D’Antonio *et al.* 2000; Ainsworth and Kauffman 2013). These introduced herbaceous and grass communities restrict native seedling recruitment (Ainsworth and Kauffman 2010) and are even more susceptible to future fires, perpetuating a grass-fire cycle (D’Antonio and Vitousek 1992; Freifelder *et al.* 1998). It has been shown that wildfires can mediate transitions between dominance by different invasive grasses (Hughes *et al.* 1991; D’Antonio *et al.* 2001). Finally, fire-adapted grasses are moving upwards in elevation (Angelo and Daehler 2013). Taken together, these studies suggest that wildfires in French Polynesia may reduce abundances of some native plant species, while simultaneously increasing abundances of some introduced species—particularly introduced grasses—which may create fuels that increase the probability of future wildfires.

### This study

The purpose of this paper is to provide a baseline for human-caused wildfire effects on species richness, abundance, and community similarity in low-elevation, heavily-invaded plant communities in French Polynesia in the modern, post-European contact era through analysis of paired field sites. In this study, we examine vegetation changes associated with wildfires at low elevations (< 380 m) on the island of Mo’orea in the Society archipelago of French Polynesia (Fig. 1). We investigate how low-elevation plant communities differ between burned areas and nearby comparison areas that have no known recent burn history, and expect wildfire to have substantial effects on the composition and relative abundances of plant species in our study sites. Comparison areas in this study do not represent natural vegetation communities, but instead are secondary forests dominated by introduced species that have regrown in areas burned or cultivated by humans following human colonization ∼1000 yr BP (Mueller-Dombois and Fosberg 1998; Kirch 2000). Native forests in French Polynesia, which harbor most remaining endemic plant and invertebrate diversity, are primarily restricted to higher elevations (Meyer 2010), and do not naturally experience large wildfires.

**Figure 1.**
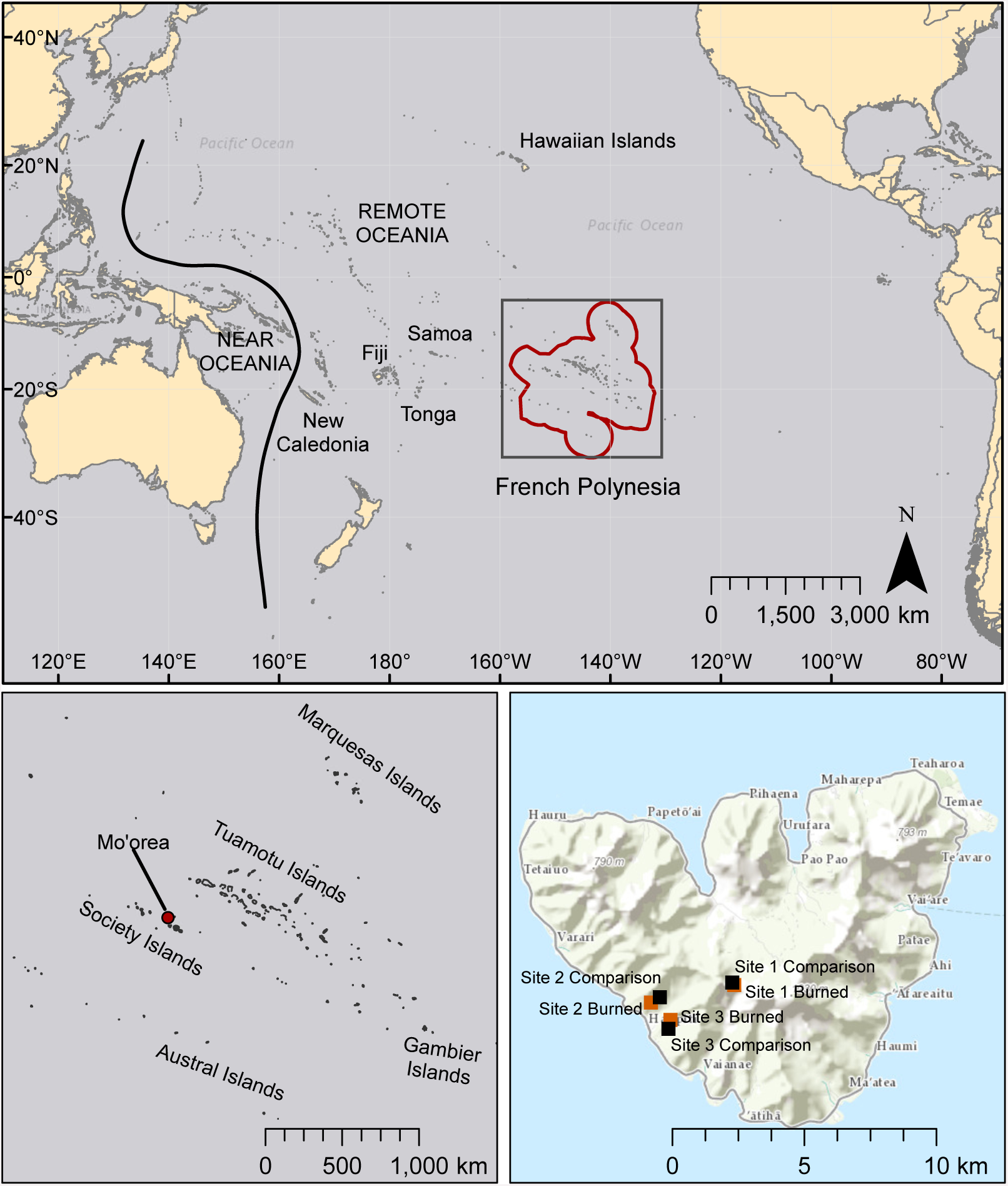
Map of the Pacific region showing French Polynesia (red outline), the island of Mo’orea, and the location of field sites. Other island chains mentioned in this study are also labeled. Basemaps are licensed through Esri (service layer credits: Esri, DeLorme, HERE, MapmyIndia, and other contributors).

Given what is known from previous observations from other islands, we hypothesize that burned areas will have a significantly different vegetation community from not recently burned or modified comparison areas. We expect that grasses and ferns, especially *Dicranopteris linearis*, will be present at higher proportions in burned areas than in comparison areas. We also expect that burned areas will contain more introduced species and higher abundances of introduced species than are present in comparison areas, but that native (especially herbaceous) vegetation will persist or recolonize burned sites (Ainsworth and Kauffman 2009; 2013). We investigate whether site or burned status (burned or comparison) is more significant in explaining species diversity and observed abundance in study areas. We examine which plant species are present through colonization, regrowth, or persistence, in burned areas. One aim of this study is to identify which of the species existing in an already highly modified plant community are likely to become more abundant or invasive after wildfire, with special attention to modern invasives and plants that are known to increase fire hazard. These results may have implications for conservation efforts in southeastern Polynesia and elsewhere in Oceania.

## Methods

We conducted research on the island of Mo’orea, French Polynesia (17°32’S, 149°50’W) during the austral-spring dry season of October and November 2013 (Fig. 1). The wetter summer season lasts from December-April, and the drier winter season lasts from May-November. Large rainfall events can occur in the mostly dry winter months, and there can be substantial interannual variation in cumulative rainfall. Weather station measurements provided by the Moorea Coral Reef Ecosystem LTER for August 2006-October 2010 report monthly average temperatures ranging from 24.5-27.2° C, and monthly cumulative rainfall between 0.8mm-716.8mm (Washburn *et al*. 2013). Winds are generally stable and dominated by southeasterly trade winds, although powerful wind events, including cyclones, occur somewhat regularly (http://mcrlter.msi.ucsb.edu/data/variable/).

### Study sites

We worked at three sites, each containing two study areas: a post-wildfire (“burned”) area that was paired with a nearby, “comparison” area (within 0.2-0.5 km) that had no recent history of fire or agricultural modification. Post-wildfire sites represented a variety of times since fire disturbance, total burned area, original vegetation communities, elevations, slopes, aspects, and microclimates. We chose to work in sites previously studied by Koehler (1999), where burned and comparison areas at a single site had the same or very similar slopes, aspects, average rainfalls, and soil types (Bonvallot *et al.* 1993), and an additional site with similar description. The seasonality and severity of the burns were not controlled for in analyses, and the post-wildfire climatic conditions are not known, each of which can influence species richness and abundance in post-wildfire communities. We clarify that this collection of sites in not meant to represent plant community succession for different-aged sites, because we did not measure or control for site-to-site variation in temperature, precipitation, and other factors important in structuring plant communities.

Study sites varied in location, time since burn, and other characteristics, but were all located on the wetter side of the island (Table 1). Site 1 was located at the Col des Trois Cocotiers (accessible via the “Three Coconuts trail”) on the saddle between Mount Tohi’e’a and Mount Mou’aroa. This wildfire originated from a small campfire at the top of the ridge (J.-Y. Meyer, *pers. comm.*) in 2004 (9 years before this survey was conducted), burning <1 ha. The site receives an average annual rainfall of 4500-5000mm, was characterized in 1993 as having an average annual temperature of 20-22.7°C (Bonvallot *et al.* 1993), and is in direct sunlight throughout the day. The slope at this site varies with a maximum of 30%. The adjacent comparison area was similar to the burned area but receives slightly more shade, and was located to the west of the burned area.

**Table 1.** Summary of characteristics for burned and comparison sites, which have no recent history of disturbance. Centers of burned areas were determined from aerial photos.

| Name of site and location | Age at time of sampling (year of burn) | Condition | Latitude/ Longitude | Elevation (m) | Size of burned area | No. species (no. observations) |
| --- | --- | --- | --- | --- | --- | --- |
| Site 1: Col des Trois Cocotiers | 9 years (2004) | burned | 17°32'49.9"S<br>149°50'32.5"W | 374 | < 1 ha | 26 (179) |
|  | (NA) | comparison | 17°32'47"S<br>149°50'35"W | 379 | (NA) | 30 (331) |
| Site 2: North of Ha'apiti | 18 years (1995) | burned | 17°33'10.39"S<br>149°52'14.63"W | 154 | ~4 ha | 18 (150) |
|  | (NA) | comparison | 17°33'4.09"S<br>149°52'3.68"W | 146 | (NA) | 21 (212) |
| Site 3: Southeast of Ha'apiti | 22 years (1991) | burned | 17°33'30.44"S<br>149°51'50.48"W | 120 | 5 ha | 13 (160) |
|  | (NA) | comparison | 17°33'40.98"S<br>149°51'53.15"W | 45 | (NA) | 17 (167) |

Site 2 is also located on the southwest side of the island, north of the town of Ha’apiti. According to Koehler (1999), the fire occurred in 1995 (18 years before survey), starting in a noni (*Morinda citrifolia*, or “*nono*” in Tahitian) fruit plantation and traveling up the slope of a west-facing ridge for approximately 1 km. The width of the burned area was ∼300 m, with a total area of ∼4 ha. The slope of this site averaged approximately 25% slope, and the ridge can receive direct light for the entire day. The average annual precipitation is 1720-2000 mm, with an average annual temperature of >22.7° C (Bonvallot *et al.* 1993). The adjacent comparison area was located to the northeast of the burned area.

Site 3 was located on the southwest side of the island, southeast of Ha’apiti. According to Koehler (1999), the wildfire occurred in 1991 (22 years before this survey was conducted), and started as a pile of burning waste in a cemetery on the outskirts of Ha’apiti. The fire traveled along a west-facing ridge for approximately 1.5 km. The total burned area was ∼5 ha, nowhere exceeding 250 m in width. The slope is ∼25% along the entire length of the burned site (Koehler 1999). The ridge receives direct sunlight for the majority of the day, the average annual precipitation at the site is 2000-2350mm, and the average annual temperature is >22.7°C (Bonvallot *et al.* 1993). The adjacent comparison area was located to the southeast of the burn. Center points of all burned and comparison areas, and total area burned were estimated using Google Earth (Google 2013).

### Field surveys

Each of the paired study areas at three sites (Sites 1, 2, and 3) comprises a post-wildfire burned area and an adjacent, unburned comparison area, for six total study areas. Five transect lines were placed in each study area, for a total of 30 transects. Transect lines were placed approximately 4 meters apart in comparison areas and in burn sites where the shape of the post-fire area allowed. For long burn-scar sites, transects were placed slightly closer to one another or end-to-end. Vegetation was sampled at 4m intervals along the length of each transect (starting at the zero point, for a total of 13 samples per transect and *n* = 390 total data points) at all sites along points on a 50m transect tape, using a method modified from the Line Interception Method (or “line-intercept method,” Canfield 1941).

For each burned and comparison area, all individuals of vascular plant species touching and directly above a vertical measuring post placed at 4m intervals were recorded. Lichens and mosses were not included in this study. Individual plants were counted on transect points (as “observed abundance”) and identified to the species level in the field if possible, and photographs were taken for later verification. For plants that could not be field-identified, samples were collected for examination at the University of California, Berkeley Gump Station using online and printed plant identification keys (Moorea Digital Flora Project, Murdock 1999; Whistler 1995). Samples and photographs were verified through expert opinion (of J.-Y. Meyer, R. Taputuarai, and J. Nitta). Voucher specimens collected from each site have been deposited in the University Herbarium (UC) at the University of California, Berkeley. Field surveys were conducted under the authorization of a University of California Berkeley permit associated with undergraduate educational course (ESPM C107) from the Government of French Polynesia.

### Analyses

Plant species were categorized by origin as native or introduced, and within the category of introduced species, a distinction was made between “Polynesian introduction” (prior to European contact) and “modern introduction” (post European contact) for analysis (Table 2). Historical presence was assessed by consensus and expert opinions including the following: Austin (1991), Butaud *et al.* (2008), Butaud (2010), Florence (1997, 2004), Florence *et al.* (2007), McCormack (2007), Meyer (2004), J.-Y. Meyer (*pers. comm*.), J. Nitta (*pers. comm*.), Prebble (2008), and Whistler (1996, 2009).

**Table 2.**
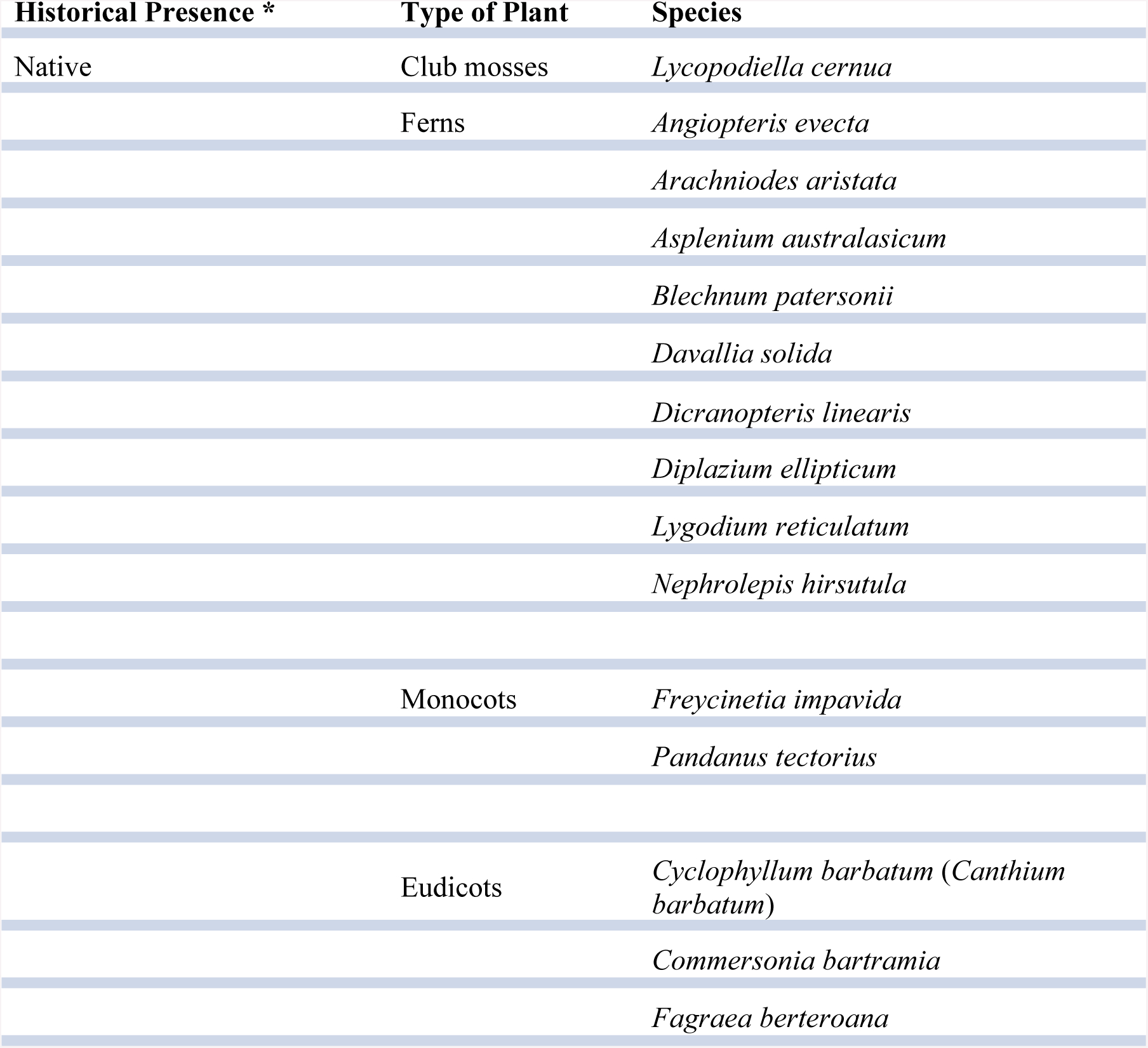
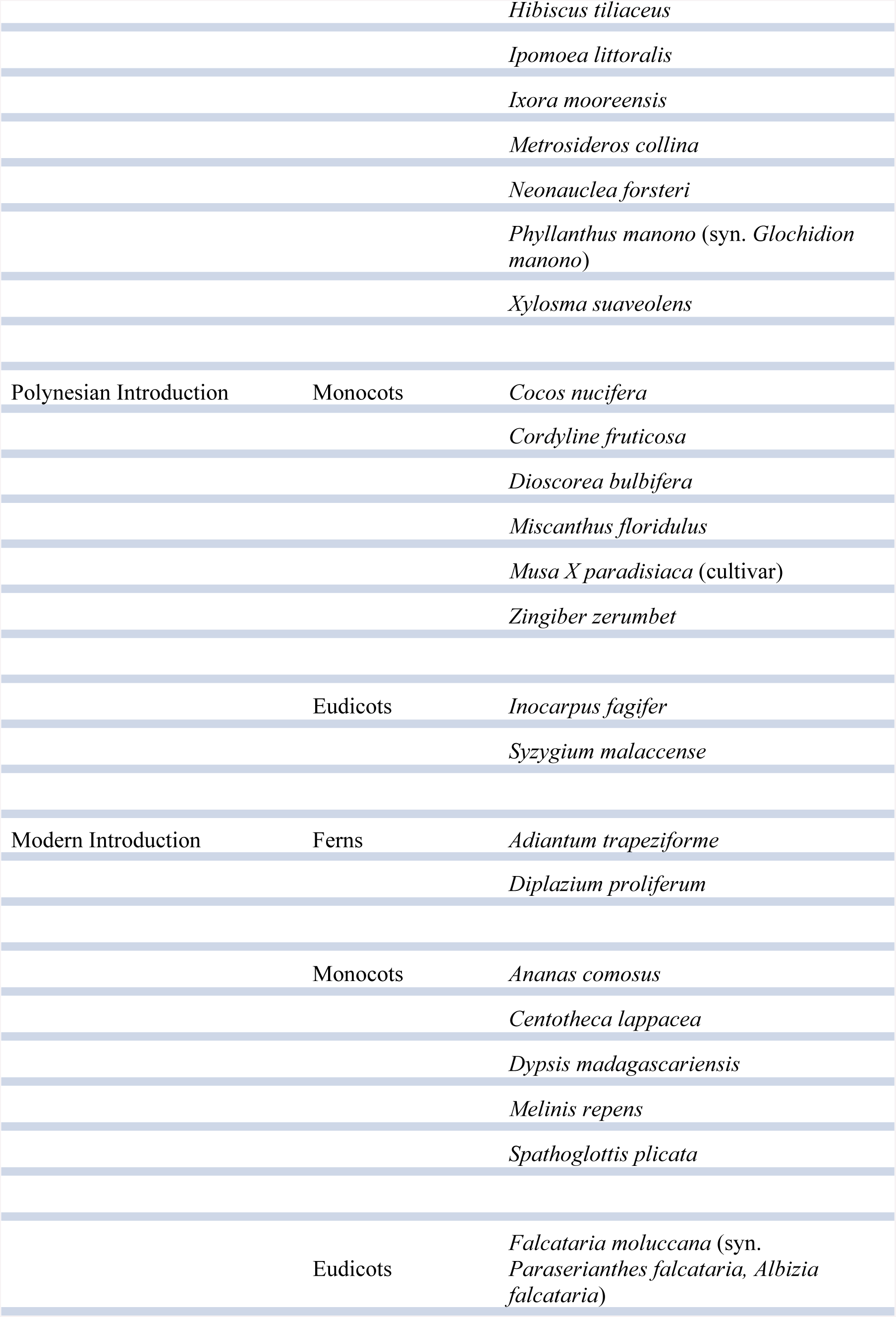
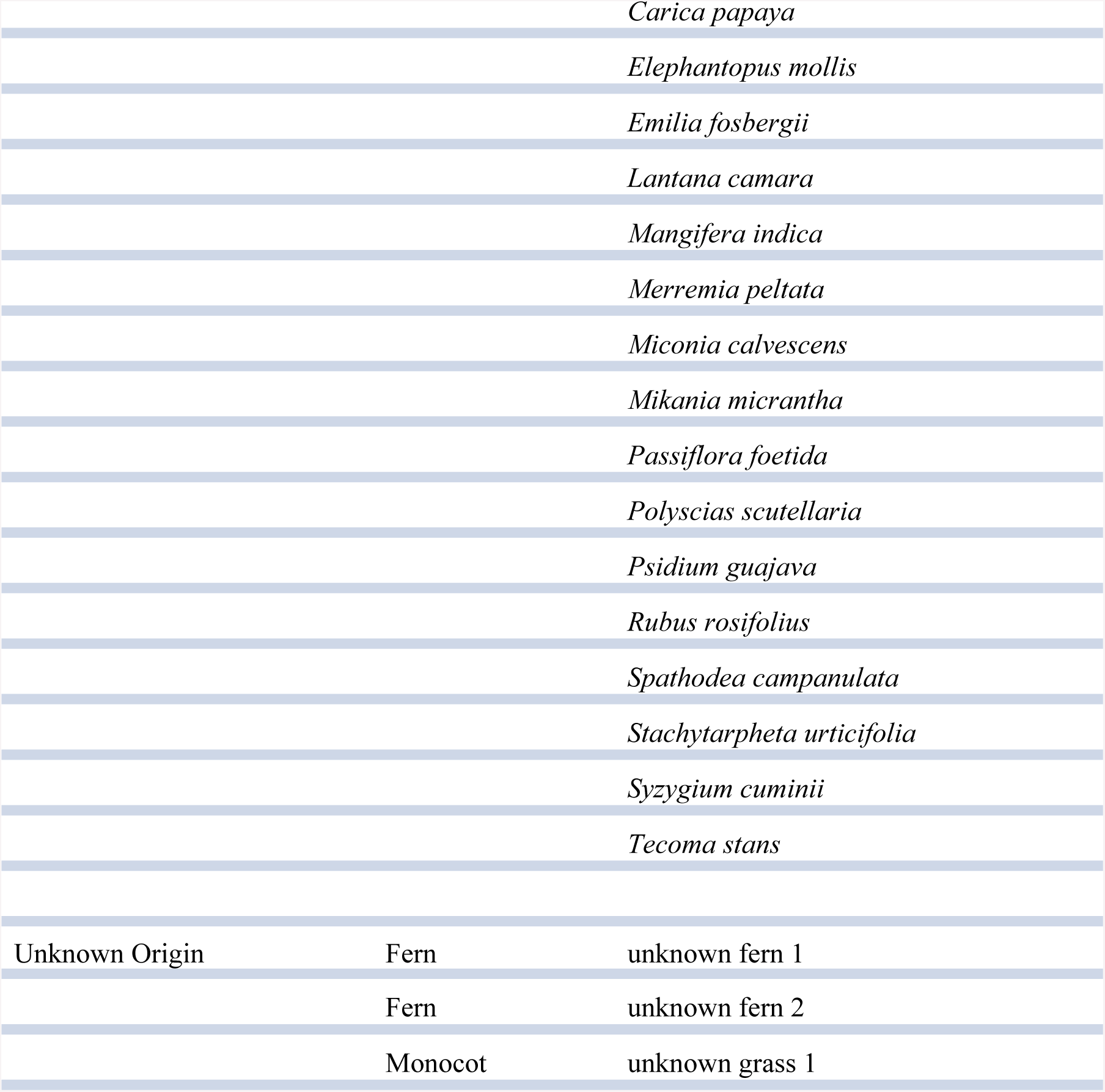
Plant species recorded at the three study sites, categorized by broad taxonomic groups. Species are also categorized by their historical presence in the Society Islands: e.g., native, introduced by humans before 1767 C.E. (“Polynesian introductions”), or introduced by humans since 1767 C.E. (“modern introductions”). Lichens and mosses were not included in this study. Historical presence was assessed by consensus and expert opinions including the following references: Austin 1991, Butaud et al. 2008, Butaud 2010, Florence 2004, Florence 1997, Florence et al. 2007, McCormack 2007, Meyer 2004, J.-Y. Meyer (*pers. comm*.), J. Nitta (*pers. comm.*), Prebble 2008, Whistler 1996, and Whistler 2009.

Line-intercept transect data were used to calculate relative frequencies of species’ abundances for burned and comparison areas at each of the three study sites. Observed abundances were calculated as total number of individuals in each species pooled across all five transects in one study area. Total species richness was calculated as number of species encountered on all transects in each area. Plants were organized for analysis by presence in burned areas, comparison areas, or both, and graphed by category for further examination. Differences between species richness by category of origin was assessed with Pearson’s Chi-squared tests for proportions, with the null hypothesis that the burned areas have the same proportions of species richness by category of origin as the comparison areas. Similar calculations were made for observed abundances of plants by category of origin.

To test the relative importance of site (SITE) and burned or comparison status (STATUS) in determining the plant community in an area, we constructed separate sets of Poisson-distributed generalized linear models (GLMs) for species richness, and for observed abundances at each transect point (Poisson and Gaussian models were run in each case and compared to each other with AIC and log likelihood values). We included an interaction effect of SITE:STATUS in each model. Akaike’s Information Criterion adjusted for small sample sizes (AIC*c*) was applied to each model set for model selection.

For plant data in each survey area, we performed a two-dimensional non-metric multi-dimensional scaling (NMDS) using a Bray-Curtis dissimilarity matrix to assess community-level differences among sites and between burned and comparison areas, using individual transects as data points. Bray-Curtis NMDS analyses evaluate community composition and structure. We employed Wisconsin double standardization (using R package “vegan” “metaMDS” analysis, Oksanen *et al.* 2018). Results of the NMDS were evaluated using a permutational multivariate analysis of variance (“adonis” in R software package “vegan”) modeling the NMDS against burned or comparison status of study areas. A statistically significant value of the “adonis” *F*-test would indicate that the burned areas are different from the comparison areas in terms of community composition, measured in squared deviation from the centroid of each group in ordination space.

All analyses were conducted in “R” version 3.1.3 (R Core Team 2015), with additional packages “vegan” version 2.5-2 (Oksanen *et al.* 2018), “MASS” version 7.3-34 (Venables and Ripley 2002), and “AICcmodavg” version 2.1-1 (Mazerolle 2017).

## Results

### Fire effects on species composition

Wildfire at all surveyed sites had large effects on the species composition of the resulting communities, in contrast to comparison areas. Species that were only present in comparison areas (23 species), outnumbered those that were only present in burned areas (11 species), but showed comparable richness to species that were present in both burned and comparison areas (23 species) (Fig. 2). Some of the most common species present only in comparison areas included the native fern *Angiopteris evecta*, the Polynesian-introduced ginger *Zingiber zerumbet*, the modern introduction *Miconia calvescens*, and the coconut *Cocos nucifera*. In burned areas, some of the most common species were the native fern *Dicranopteris linearis* and the native tree *Metrosideros collina*. Species found in all sites included the native tree *Hibiscus tiliaceus* and the modern introduction *Paraserianthes falcataria.*

**Figure 2.**
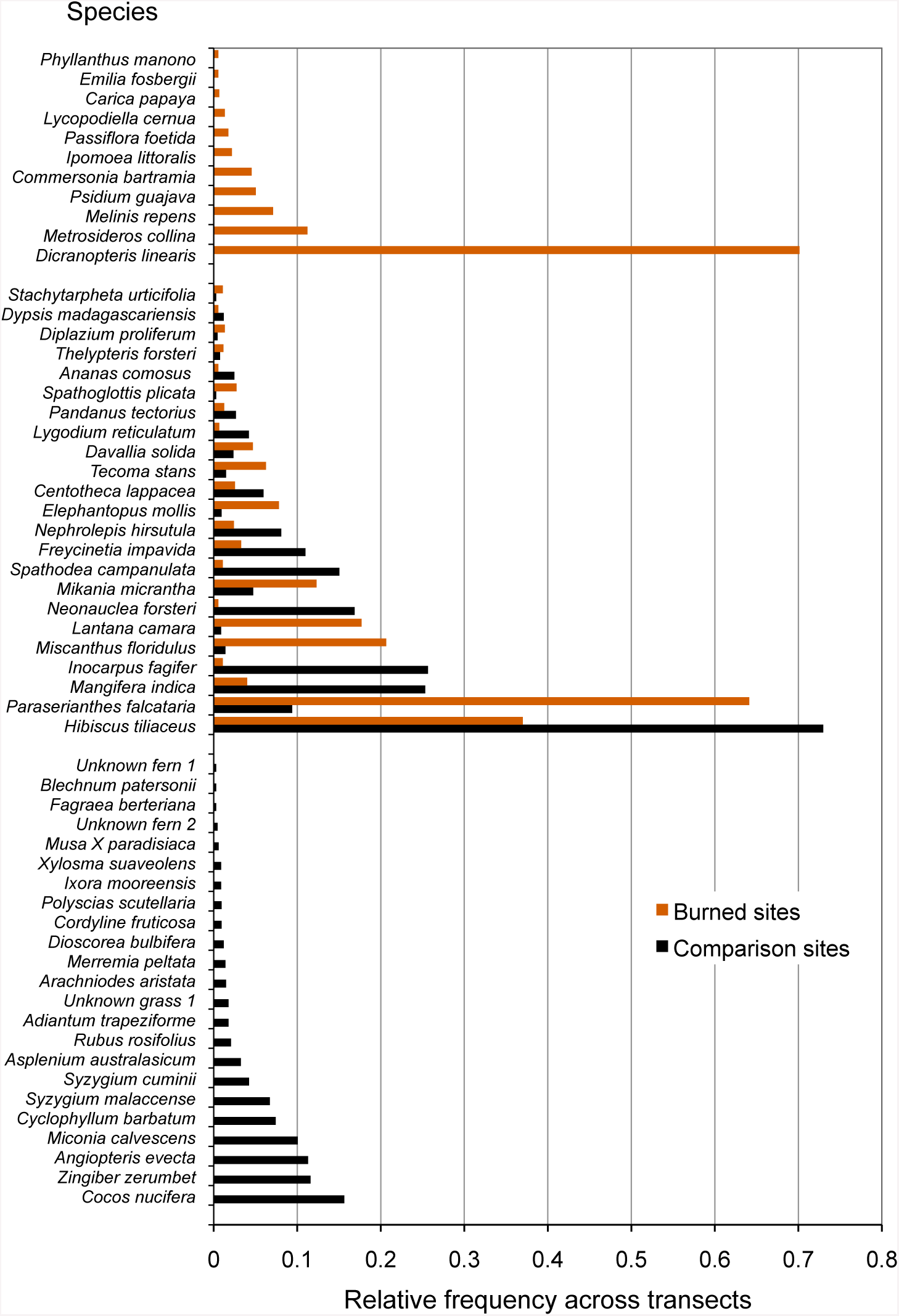
Species recorded in burned and comparison areas, arranged by relative abundance. Data represent all individuals recorded at all sites, and are grouped by presence in burned areas, comparison areas, or both.

### Fire effects on species richness and abundance

Burned and comparison comparison sites contained a total of 57 vascular plant species (one club moss, 13 ferns, 14 monocots, and 29 eudicots; Table 2). Of the vascular plants, 54 were identified to the species level, and three unknown species were given unique identifiers for analysis. At comparison sites, 46 species were found in total, with 34.8% native species, 17.4% Polynesian introduced species, and 41.3% modern introduced species (with three unidentified species of unknown origin). By comparison, 34 species were found in burned sites, 38.2% of which were natives, 5.9% Polynesian, and 50.0% modern introductions (Fig. 3-Fig. 4; raw data visualized in Supplementary Material). Observed abundances varied on a site-to-site basis, with wildfire increasing observed abundances at two of three sites, and sharply decreasing observed abundances at Site 2.

**Figure 3.**
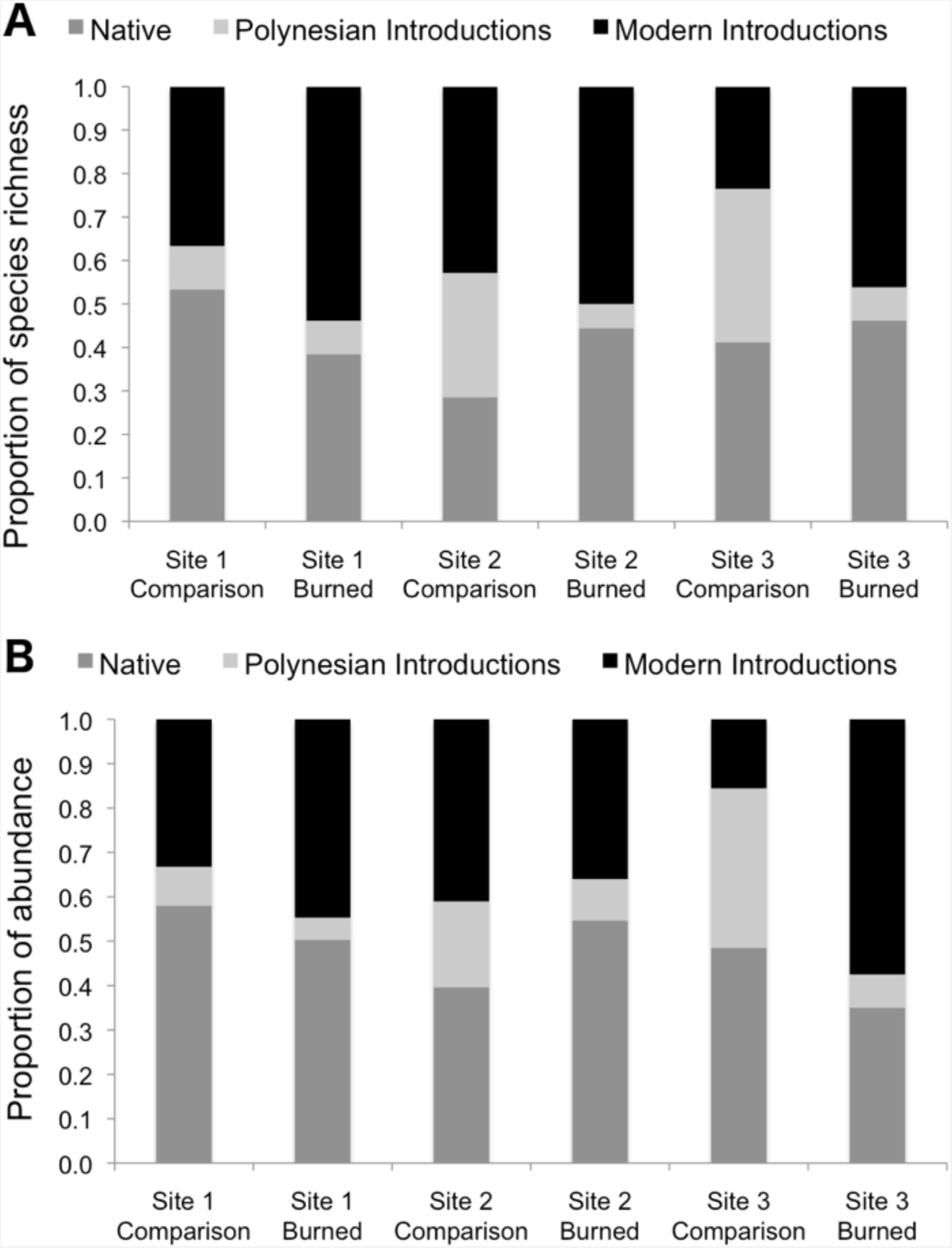
Species richness (A) and observed abundances of species (B) shown as proportions by category of origin, measured at sampling points in burned and comparison areas at 3 sites. Data represent individuals measured at 13 points each along 5 transects in burned and comparison areas for each of 3 sites (n = 390 data points). Plants are grouped into native, Polynesian introduction, and modern introduction categories.

**Figure 4.**
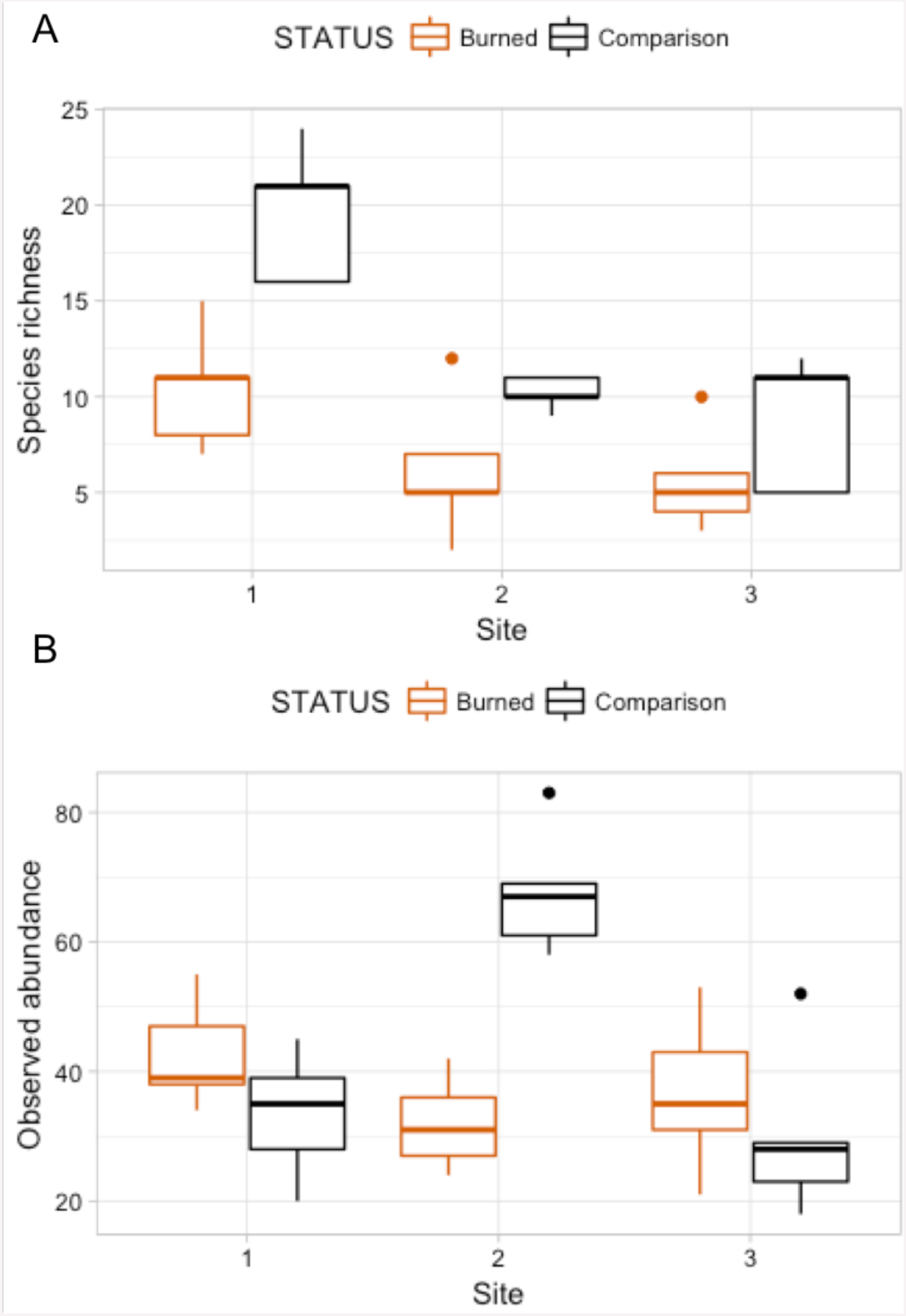
Box plots comparing species richness and observed abundances across burned and comparison areas (“Status”) at three sites.

A Poisson model of species richness ∼ SITE + STATUS had the most explanatory power (evidence ratio compared to next best model = 16.42) (Table 3A). We found that comparison areas had greater species richness than burned areas **(*P <* 0.001)**; Site 1 had the greatest species richness (Table 4). GLM analysis of observed abundances (Table 3B) produced one model with explanatory power (abundance = SITE + STATUS + SITE:STATUS). Fixed effect parameter estimates (Table 5) show that “comparison” status had higher abundance generally than burned areas **(*P* = 0.079)**, though the effect size was relatively small (0.152). As was the case for the species richness GLM, the high point estimate and statistical significance of the intercept in each case indicates that more nuanced models with higher explanatory power could be developed.

**Table 3.** Model selection for species richness and for observed abundance. Models are shown in rank order of AICc support. Here, k = number of parameters in model; AICc = Akaike’s Information Criterion value corrected for small sample sizes; ΔAICc = difference of AICc value compared to the next best-supported model; wi = AICc weight (a measure of strength of evidence for each model); Cumulative w = total model AICc weights for best models in rank order; and LL = Log-Likelihood.

| Ranked models for $y = \text{SPECIES}$ | | | | | | |
| --- | --- | --- | --- | --- | --- | --- |
| Model family = Poisson | $k$ | AICc | $\Delta\text{AICc}$ | $w_i$ | Cumulative $w$ | LL |
| $y = \text{SITE} + \text{STATUS}$ | 4 | 156.90 | 0 | 0.94 | 0.94 | -73.65 |
| $y = \text{SITE} + \text{STATUS} + \text{SITE}:\text{STATUS}$ | 6 | 162.49 | 5.60 | 0.06 | 1 | -73.42 |
| $y = \text{SITE}$ | 3 | 176.61 | 19.72 | 0 | 1 | -84.85 |
| $y = \text{STATUS}$ | 2 | 185.48 | 28.58 | 0 | 1 | -90.52 |
| $y = 1$ | 1 | 205.57 | 48.67 | 0 | 1 | -101.71 |

(B)
| Ranked models for $y = \text{ABUNDANCE}$ | | | | | | |
| --- | --- | --- | --- | --- | --- | --- |
| Model family = Poisson | $k$ | AICc | $\Delta\text{AICc}$ | $w_i$ | Cumulative $w$ | LL |
| $y = \text{SITE} + \text{STATUS} + \text{SITE}:\text{STATUS}$ | 7 | 237.02 | 0 | 1 | 1 | -108.96 |
| $y = \text{SITE}$ | 4 | 253.67 | 16.66 | 0 | 1 | -122.04 |
| $y = \text{SITE} + \text{STATUS}$ | 5 | 254.89 | 17.88 | 0 | 1 | -121.2 |
| $y = 1$ | 2 | 255 | 17.98 | 0 | 1 | -125.28 |
| $y = \text{STATUS}$ | 3 | 256.13 | 19.11 | 0 | 1 | -124.6 |

**Table 4.** Fixed effect parameter estimates from GLM analysis of total species richness, from the top AICc-selected model. Significant parameters are printed in **bold**.

| Parameter | Point Estimate | Standard error | *P*-value |
| --- | --- | --- | --- |
| **Intercept** | **2.3937** | **0.1113** | **< 0.001** |
| **SITE THREE** | **-0.7340** | **0.1434** | **< 0.001** |
| **SITE TWO** | **-0.6039** | **0.1373** | **< 0.001** |
| **STATUS (Comparison)** | **0.5532** | **0.1191** | **< 0.001** |

**Table 5.** Fixed effect parameter estimates from GLM analysis of observed abundance, from the top AICc-selected model. Significant parameters are printed in **bold**.

| Parameter | Point Estimate | Standard error | *P*-value |
| --- | --- | --- | --- |
| **Intercept** | **3.68819** | **0.0973** | **< 0.001** |
| SITE | -0.06368 | 0.0410 | 0.138 |
| **STATUS (Comparison)** | **0.15209** | **0.0831** | **0.079** |
| SITE:STATUS (Comparison) | 0.04208 | 0.0707 | 0.570 |

### Fire effects on proportions of native and introduced species

Species richness was compared between study areas by category of origin (Fig. 3). Pearson’s Chi-squared tests for proportions show no differences between proportions of species in the native, Polynesian introduction, and modern introduction categories for burned and comparison areas at each site (*χ*^2^ = 1.43, df = 2, *P* = 0.434 at Site 1; *χ*^2^ = 3.65, df = 2, *P* = 0.161 at Site 2; and *χ*^2^ = 3.58, df = 2, *P* = 0.167 at Site 3). When modern and Polynesian introductions are grouped into one “introduced species” category, this result does not change, that is, there are no statistically significant differences between proportions of native and introduced species by native versus introduced categories (*P*-values on *χ*^2^ tests are all > 0.2).

Proportional abundances of all individuals were also compared across burned and comparison areas in three sites by category of origin (Fig. 3). Pearson’s Chi-squared tests show statistically significant differences between proportions of abundances of individuals in the native, Polynesian introduction, and modern introduction categories for burned and comparison areas at each site (*χ*^2^ = 7.523, df = 2, ***P* = 0.023** for Site 1; *χ*^2^ = 10.697, df = 2, ***P* = 0.005** for Site 2; and *χ*^2^ = 76.705, df = 2, ***P* < 10^−5^** for Site 3). When modern and Polynesian introductions are grouped into one “introduced species” category, results are still statistically significant (*P* = 0.09 for Site 1**; *P* = 0.005** for Site 2; and ***P* = 0.01** for Site 3). Wildfire significantly increased the proportional abundance of individuals in the “introduced species” category at Sites 1 and 3, but decreased the proportional abundance of introduced species at Site 2.

### Fire effects on plant communities

Plant species composition and abundances by transect were analyzed using two-dimensional NMDS, which converged in 20 iterations, for a calculated stress = 0.160 (considered an acceptable fit), Procrustes root mean squared error = 1.64e-05, and maximum residuals = 2.98e-05. NMDS results were plotted with individual points representing species richness and observed abundances of a single transect (Fig. 5). There was no overlap in the points between the burned and comparison areas, visually indicating large differences between burned and comparison communities in ordination space. Points for each of the comparison areas were highly clustered and did not overlap with points from other sites, indicating substantial differences in communities between different (not recently burned) comparison areas. In contrast, points for the burned areas are highly mixed and essentially indistinct from each other, and within-site spread of points in NMDS-ordination space is larger for the burned areas than for the comparison areas. The “adonis” permutational multivariate analysis of variance carried out on the NMDS produced test statistic values of *F*_1,28_=14.734 and *R*^*2*^ = 0.345, with ***P* < 0.001**, meaning that compositional differences between groups (in this case burned and comparison areas) are highly statistically significant.

**Figure 5.**
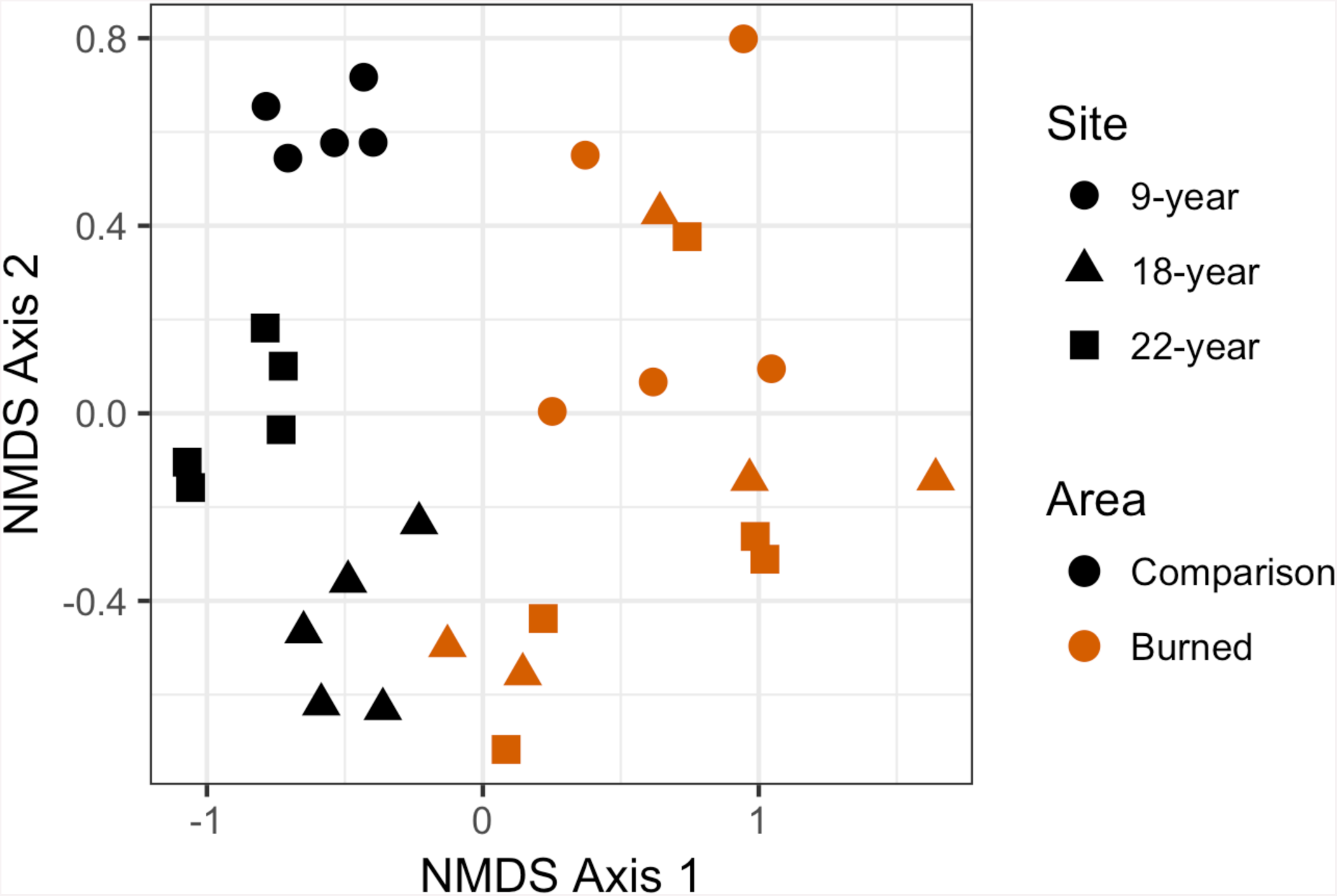
Non-metric multidimensional scaling (NMDS) plot showing differences in plant community composition between transects from each site. Points that are closer together have a higher degree of similarity. Each point represents a single transect; burned area data are represented in red, and comparison plot data are represented in black. Post-fire areas have larger and more overlapping ranges in this parameter space than do comparison areas.

## Discussion

Even in an already heavily impacted plant community, wildfire in low-elevation Pacific island vegetation creates communities that are very similar between sites, whereas comparison sites with no recent history of disturbance remain relatively distinct from one another. Wildfire does not appear to change the proportion of species in a community by their category of origin (*i.e*. native, Polynesian introduction, or modern introduction). Counter to our original hypothesis, we did not find more introduced species in burned areas than comparison areas. Observed abundances were generally higher in comparison areas than burned areas, but the effect of wildfire was highly variable between sites, and wildfire greatly lowered observed abundance at one of three sites (Site 2), possibly due to post-wildfire conditions for regeneration or other unmeasured variables. Our NMDS results (followed by multivariate variance analysis) also show significant differences in species composition of burned and comparison areas.

These results taken together provide evidence that wildfire impacts on the native community of plants vary from site to site, and may be related to microclimate or other environmental drivers affecting the original vegetation community (Thaxton and Platt 2006; Hoffman *et al.* 2011), or properties of the fire events themselves (such as temperatures, intensity, heat flux, residence time; Frost and Robertson 1985; Thaxton and Platt 2006), and post-fire climatic conditions (Frost and Robertson 1985) that were not measured in this study.

Our results suggest that there is a significant component of the low-elevation plant community that is directly benefitting from wildfire activity and are present only in burned areas. We found, as previous studies have, that the native fern *Dicranopteris linearis* (Gleicheniaceae) colonized burned areas, consistent with observations of the pyrophytic *Dicranopteris* savanna noted by other investigators (Papy 1954; Mueller-Dombois and Fosberg 1998; Meyer 2017). We did not find higher presence of grasses in general, as has been noted in Hawai‘i (Hughes *et al.* 1991; Hughes and Vitousek 1993; D’Antonio *et al.* 2000; Ainsworth and Kauffman 2013), although we found that the non-native invasive grass *Melinis repens* (natalgrass, Poaceae) was present only in the burned sites. This species is considered invasive in both French Polynesia and Hawai’i, and the same pattern—presence in burned areas only— was reported by D’Antonio *et al*. (2011). It should be noted that all our study sites were located on the wet side of the island, and that non-native grasses are present in abundance only on the dry side of Mo’orea. A study designed to sample the dry side of Mo’orea might discover additional invasives that may be spreading through interaction with wildfire.

Among other species found only in burned areas are the native trees *Metrosideros collina* (Myrtaceae) and *Commersonia bartramia* (Malvaceae). This is consistent with observations of *M. collina* in association with either fire or *Dicranopteris* savanna (*e.g*., Papy 1954) and of *C. bartramia* in *Dicranopteris* savanna (Florence 2004; Meyer 2017). One observation of the native tree *Phyllanthus manono* (syn. *Glochidion manono*, Phyllanthaceae) in a burned area is interesting to the natural history of this species given that this genus has often been reported from disturbed contexts in Remote Oceania (Hembry 2017), but it was not noted at the time of the survey if this was an individual that had resprouted or grown from seed. Of other species found in association with fire, pineapple (*Ananas comosus*, Bromeliaceae) is a fruit crop that may have been planted following the fires, and therefore likely does not represent a “naturalized” species benefitting from fire. Papaya (*Carica papaya*, Caricaceae), on the other hand, does seem to benefit from fire in western Polynesia (Franklin 2007) although we are not aware of evidence that it is naturalized in southeastern Polynesia where this study took place (*e.g*., Sykes 2016).

Of the species present in both burned and comparison areas, the native tree *Hibiscus tiliaceus* (Malvaceae) is the most abundant, followed by a problematic modern invasive, *Paraserianthes falcataria* (Fabaceae). This species (syn. *Falcataria molluccana, Albizia falcataria*), when compared to other invasives, has been identified as having disproportionate effects on species loss in tropical systems it invades (Pyšek *et al.* 2012). Several native and introduced overstory trees are present in both burned and comparison sites, but are more common in the latter. These include the native *Neonauclea forsteri* (Rubiaceae), the Polynesian-introduced *Inocarpus fagifer* (Fabaceae), and the modern introductions *Spathodea campanulata* (Bignoniaceae) and *Mangifera indica* (mango; Anacardiaceae). We suggest that these have merely persisted through low-intensity fire in small numbers. Persistence of overstory trees in or at the margins of low-intensity burns may be more common, and such observations should be noted in future studies on this topic.

Of the comparison-area only plants, notable are coconut (*Cocos nucifera*, Arecaceae), which may have been planted, a host of native species which may be highly fire-intolerant, including *Ixora mooreensis* (Rubiaceae) and *Xylosma suaveolens* (Flacourtiaceae), a large number of native fern species, and an invasive species of special concern in French Polynesia, *Miconia calvescens* (Melastomataceae), which densely invades otherwise undisturbed areas (Meyer and Florence 1996).

Our results highlight four introduced and/or invasive species which may be facilitated by fire and which are of concern to land management in French Polynesia and the Pacific region. Natalgrass (*Melinis repens*), a modern introduction found only in burned sites in this study, is discussed above. *Miscanthus floridulus* (Poaceae) is much more abundant in burned than comparison sites in our study, as are *Paraserianthes falcataria* and *Lantana camara*. *Miscanthus floridulus* is a Polynesian-introduced grass which is considered a noxious weed in Hawai‘i, although not in French Polynesia, and is considered native on Guam and the northern Mariana Islands, where it is a dominant component of fire-prone anthropogenic savannas (Minton 2006). Although ranked only sixth in abundance in burned sites, *Lantana camara* is a modern introduction that is probably of higher concern than some of the more abundant species. Fire-*Lantana* interactions, including the possibility of a “fire cycle” or positive feedback between the two, are of special concern and have been reported from continental tropical areas (Fensham *et al.* 1994; Duggin and Gentle 1998; Hiremath and Sundaram 2005; Berry *et al.* 2011). The flammability and other fire adaptations of this species compared to other vegetation in this system are not known. *Lantana camara* is bird-dispersed in French Polynesia (Spotswood *et al.* 2012), and so has the potential to increase its range rapidly through seed dispersal. *Lantana camara* is designated by the French Polynesian government as one of the most serious invasive plants in the country (Meyer and Butaud 2007).

The comparison sites, like most low-elevation plant communities on Mo’orea and elsewhere in French Polynesia, are secondary forest or shrub communities heavily invaded by, and in many cases dominated by, non-native species of angiosperms. These heavily invaded plant communities have developed over a land use history that includes human clearing of land for agriculture and timber, the cultivation of agroforests by Polynesians before European contact, the extinction of native birds as seed dispersers, the extirpation of seabird colonies and their nutrient inputs, and the introduction of ungulates (especially pigs; *Sus scrofa*) (Kirch 2000; Meyer 2004; Steadman 2006; Dotte-Sarout and Kahn 2017). There is no evidence to suggest that these heavily transformed vegetation communities at low elevations on Mo’orea have a natural fire regime of the sort which occurs in continental regions. Fire in this context should be thought of as an anthropogenic disturbance that has profound effects on these primarily non-native vegetation communities. We found that in this context, fire facilitates a subset of the native flora (such as *Dicranopteris linearis* and *Metrosideros collina*) but also facilitates several of the most destructive invasive plants in French Polynesia (*Paraserianthes falcataria*, and *Lantana camara*), and two introduced grasses considered invasive in Hawai‘i (*Melinis repens, Miscanthus floridulus*), one of which is listed as a noxious weed by the state of Hawai‘i (*Miscanthus floridulus*).

## Conclusions

Our results demonstrate that wildfire on Mo’orea causes significant changes in plant communities, increasing similarity of communities between sites, increasing abundances of introduced species in burned areas, and changing community composition by facilitating colonization by *Dicranopteris linearis* as well as non-native species that have known positive fire feedback interactions. These changes are present in all post-wildfire sites (including the oldest site of 22 years). Long-term changes to the vegetation community that promote non-native and fire-prone species could be especially harmful on small tropical islands because they are more susceptible to invasion, and the plant diversity is already relatively low (Elton 1958; Vitousek *et al.* 1996).

Pacific islands invasives are comparatively understudied (Pyšek *et al.* 2012), and disturbance ecology has not been studied in many Pacific island groups. Our results may be generalizable to other islands both within the Society Islands chain and elsewhere in French Polynesia and southeastern Polynesia, where the original vegetation communities are similar to those on Mo’orea, and the invasive species available to colonize post-wildfire sites come from the same pool of available species. These results are important in light of frequent recent reports of wildfire events in local news media from multiple islands in French Polynesia, and because both wildfire frequency and total burned area per year may increase in future years with climate change, presence of invasive species, increasing populations that rely on fire for agricultural and waste management, and other changes in human land use.

## Acknowledgements

We thank the instructors of the Biology and Geomorphology of Tropical Islands, G. Roderick, B. Mishler, V. Resh, J. Stillman, and S. Carlson, as well as graduate student instructors C. DiVittorio, J. Hopper, and L. Dougherty for project guidance; J.-Y. Meyer, R. Taputuarai, and J. Nitta for plant identifications; D.S. Godwin for comments and graphing code; and G. Simpson and R.B. de Andrade for comments. We especially thank C. Trauernicht for extensive comments and suggested analyses. We thank V. Brotherson and Gump Research Station for providing logistical support, field sites and facilities, and P. Frogier and the Délégation à la Recherche for providing permits to CAW, EAN, and DHH. We also thank K. Hartfield and the Arizona Remote Sensing Center for mapping support.

## References

Ainsworth A, Kauffman JB (2009) Response of native Hawaiian woody species to lava-ignited wildfires in tropical forests and shrublands. Plant Ecology 201, 197–209.

Ainsworth A, Kauffman JB (2010) Interactions of fire and nonnative species across an elevation/plant community gradient in Hawaii Volcanoes National Park. Biotropica 42, 647–655.

Ainsworth A, Kauffman JB (2013) Effects of repeated fires on native plant community development at Hawaii Volcanoes National Park. International Journal of Wildland Fire 22, 1044–1054.

Angelo CL, Daehler CC (2013) Upward expansion of fire-adapted grasses along a warming tropical elevation gradient. Ecography 36, 551–559.

Athens JS, Ward JV (2004) Holocene vegetation, savanna origins and human settlement of Guam. Records-Australian Museum 19, 15–30.

Austin DF (1991) *Ipomoea littoralis* (Convolvulaceae)—taxonomy, distribution, and ethnobotany. Economic Botany 45, 251–256.

Berry ZC, Wevill K, Curran TJ (2011) The invasive weed *Lantana camara* increases fire risk in dry rainforest by altering fuel beds. Weed Research 51, 525–533.

Bonvallot J, Dupon J-F, Vigneron E, Gay J-C, Morhange C, Ollier C, Peugniez G, Reitel B, Yon-Cassat F (1993) ‘Atlas de la Polynésie française.’ (Éditions de l’ORSTOM: Paris, France)

Butaud J-F, Gérard J, Guibal D (2008) ‘Guide des arbres de Polynésie française: bois et utilisations.’ (Éditions Au vent des îles: Pirae, French Polynesia)

Butaud J-F (2010) ‘Gambier: guide floristique.’ (Direction de l’Environnement, Government of French Polynesia: Papeete) Available online: http://www.environnement.pf/spip.php?article125

Canfield RH (1941) Application of the line interception method in sampling range vegetation. Journal of Forestry 39, 388–394.

Cochrane MA (2003) Fire ecology for rainforests. Nature 421, 913–919.

D’Antonio CM, Hughes RF, Tunison, JT (2011). Long-term impacts of invasive grasses and subsequent fire in seasonally dry Hawaiian woodlands. Ecological Applications 21, 1617–1628.

D’Antonio CM, Hughes RF, Vitousek PM (2001) Factors influencing dynamics of two invasive C_4_ grasses in seasonally dry Hawaiian woodlands. Ecology 82, 89–104.

D’Antonio CM, Tunison JT, Loh RK (2000) Variation in the impact of exotic grasses on native plant composition in relation to fire across an elevation gradient in Hawaii. Austral Ecology 25, 507–522.

D’Antonio CM, Vitousek PM (1992) Biological invasions by exotic grasses, the grass/fire cycle, and global change. Annual Review of Ecology and Systematics 23, 63–87.

Davis MA, Grime JP, Thompson K (2000) Fluctuating resources in plant communities: A general theory of invasibility. Journal of Ecology 88, 528–534

Dodson JR, Intoh M (1999) Prehistory and palaeoecology of Yap, federated states of Micronesia. Quaternary International 59, 17–26.

Dotte-Sarout E, Kahn JG (2017) Ancient woodlands of Polynesia: a pilot anthracological study on Maupiti Island, French Polynesia. Quaternary International 457, 6–28.

Duggin JA, Gentle CB (1998) Experimental evidence on the importance of disturbance intensity for invasion of *Lantana camara* L. in dry rainforest-open forest ecotones in north-eastern NSW, Australia. Forest Ecology and Management 109, 279–292.

Elmqvist T, Wall M, Berggren AL, Blix L, Fritioff Å, Rinman U (2002) Tropical forest reorganization after cyclone and fire disturbance in Samoa: remnant trees as biological legacies. Conservation Ecology 5, 10.

Ellsworth LM, Litton CM, Dale AP, Miura T (2014) Invasive grasses change landscape structure and fire behaviour in Hawaii. Applied Vegetation Science 17, 680–689.

Elton CS (1958) ‘The ecology of invasions by animals and plants.’ (Methuen: London, UK)

Fensham RJ, Fairfax RJ, Cannell RJ (1994) The invasion of *Lantana camara* L. in Forty Mile Scrub National Park, north Queensland. Australian Journal of Ecology 19, 297–305.

Florence J (1997) ‘Flore de la Polynésie française, vol. 1.’ (Éditions de l’ORSTOM: Paris, France)

Florence J (2004) ‘Flore de la Polynésie française, vol. 2.’ (IRD Éditions and Muséum national d’histoire naturelle: Paris, France)

Florence J, Chevillotte H, Ollier C, Meyer J-Y (2007) Base de données botaniques Nadeaud de l’Herbier de la Polynésie française (PAP). Available online: http://www.herbier-tahiti.pf

Franklin J (2007) Recovery from clearing, cyclone and fire in rain forests of Tonga, South Pacific: Vegetation dynamics 1995-2005. Austral Ecology 32, 789–797.

Freifelder RR, Vitousek PM, and D’Antonio CM (1998) Microclimate change and effect on fire following forest-grass conversion in seasonally dry tropical woodland. Biotropica 30, 286–297.

Frost PGH, Robertson F (1985) Fire the Ecological Effects of Fire in Savannas. In ‘Determinants of Tropical Savannas’ (Eds Walker TS, Walker BH) pp. 93–140. (Published 1987 by IRL Press, Oxford, The International Union of Biological Sciences IUBS Monograph Series, No. 3)

Google (2013) Google Earth (Version 7). Available online: http://www.google.com/earth/

Hembry DH (2017) Phyllantheae-*Epicephala* mutualistic interactions on oceanic islands in the Pacific. In ‘Obligate pollination mutualism’. (Eds M Kato, A Kawakita) pp. 221–248. (Springer Japan: Tokyo, Japan)

Hiremath AJ, Sundaram B (2005) The fire-lantana cycle hypothesis in Indian forests. Conservation and Society 3, 26–42.

Hjerpe J, Hedenås H, Elmqvist T (2001) Tropical rain forest recovery from cyclone damage and fire in Samoa. Biotropica 33, 249–59.

Hoffmann WA, Jaconis SY, Mckinley KL, Geiger EL, Gotsch SG, Franco AC (2012) Fuels or microclimate? Understanding the drivers of fire feedbacks at savanna–forest boundaries. Austral Ecology 37, 634–643.

Hughes F, Vitousek PM (1993) Barriers to shrub reestablishment following fire in the seasonal submontane zone of Hawai’i. Oecologia 93, 557–563.

Hughes F, Vitousek PM, Tuniston T (1991) Alien grass invasion and fire in the seasonal submontane zone of Hawai’i. Ecology 72, 743–747.

Hunter-Anderson RL (2009) Savanna anthropogenesis in the Mariana Islands, *Micronesia: re-interpreting the palaeoenvironmental data*. Archaeology in Oceania 44, 125–141.

Kahn JG, Nickelsen C, Stevenson J, Porch N, Dotte-Sarout E, Christensen CC, May L, Athens JS, Kirch PV (2015) Mid-to late Holocene landscape change and anthropogenic transformations on Mo ‘orea, Society Islands: A multi-proxy approach. The Holocene 25, 333–347.

Kato-Noguchi H, Saito Y, Suenaga K (2012) Involvement of allelopathy in the establishment of pure colony of *Dicranopteris linearis*. Plant Ecology 213, 1937–1944.

Keppel G, Morrison C, Meyer J-Y, Boehmer HJ (2014) Isolated and vulnerable: the history and future of Pacific Island terrestrial biodiversity. Pacific Conservation Biology 20, 136–145.

Kirch PV (2000) ‘On the road of the winds: an archaeological history of the Pacific islands before European contact.’ (University of California Press: Berkeley, CA, USA)

Koehler TB (1999) Plant recovery on fire scars on Moorea, French Polynesia. University of California, Berkeley, Biology and Geomorphology of Tropical Islands 8, 71–79.

Mann D, Edwards J, Chase J, Beck W, Reanier R, Mass M, Finney B, Loret J (2008) Drought, vegetation change, and human history on Rapa Nui (Isla de Pascua; Easter Island). Quaternary Research 69, 16–28.

Marc J. Mazerolle (2017) AICcmodavg: Model selection and multimodel inference based on (Q)AIC(c). R package version 2.1-1.

McCormack G (2007) Cook Islands Biodiversity Database, Version 2007.2. Cook Islands Natural Heritage Trust, Rarotonga, Cook Islands. Available online: http://cookislands.bishopmuseum.org.

Meyer J-Y (2004) Threat of invasive alien plants to native flora and forest vegetation of Eastern Polynesia. Pacific Science 58, 357–375.

Meyer J-Y (2010) Montane cloud forests in remote islands of Oceania: the example of French Polynesia (South Pacific Ocean). In ‘Tropical montane cloud forests: Science for conservation and management’. (Eds LA Bruijnzeel, FN Scatena, LS Hamilton) pp. 121–129. (Cambridge University Press: Cambridge, UK)

Meyer J-Y (2017) ‘Guide des plantes indigènes et endémiques de Wallis et Futuna (’Uvea, Futuna, Alofi).’ (Éditions Au vent des îles: Pirae, French Polynesia)

Meyer J-Y, Butaud J-F (2007) ‘Les plantes envahissantes en Polynésie française: guide illustré d’identification.’ (Government of French Polynesia, Direction de l’Environment and Délégation à la Recherche: Papeete, French Polynesia).

Meyer J-Y, Florence J (1996) Tahiti’s native flora endangered by the invasion of *Miconia calvescens* (Melastomataceae). Journal of Biogeography 23, 775–781.

Minton, D. 2006. Fire, erosion, and sedimentation in the Asan-Piti watershed and War in the Pacific NHP, Guam. Tech. Rep. 150. Pacific Cooperative Studies Unit, University of Hawai‘i, Honolulu.

Mueller-Dombois D, Fosberg FR (1998) ‘Vegetation of the tropical Pacific islands.’ (Springer: Berlin, Germany).

Murdock A (1999) Moorea digital flora project. Available online: http://ucjeps.berkeley.edu/moorea/index.html.

Newman EA (2014) Fire in paradise. Wildfire Magazine, (Nov. issue) 22–26.

Nunn PD (1990) Recent environmental changes on Pacific Islands. Geographical Journal, 125–140.

Oksanen J (2011) Multivariate analysis of ecological communities in R: vegan tutorial, 43 pp.

Oksanen J, Guillaume Blanchet F, Kindt R, Legendre P, Minchin PR, O’Hara RB, Simpson GL, Solymos P, Stevens MHH, Wagner H (2018) vegan: Community Ecology Package. R package version 2.5-2. http://CRAN.R-project.org/package=vegan.

Papy HR (1954) ‘Tahiti et les îles voisines, II, la végétation des îles de la Société et de Makatea.’ (Laboratoire Forestier: Toulouse, France).

Perry, GLW, and Enright NJ (2002) Humans, fire and landscape pattern: understanding a maquis-forest complex, Mont Do, New Caledonia, using a spatial “state-and-transition” model. Journal of Biogeography 29, 1143–1158.

Prebble M (2008) No fruit on that beautiful shore: What plants were introduced to the subtropical Polynesian islands prior to European contact? In ‘Islands of inquiry: Colonisation, seafaring and the archaeology of maritime landscapes’. (Eds G Clark, F Leach, S O’Connor) pp. 227–251. (Terra Australis 29). (ANU ePress: Canberra, Australia)

Pyšek P, Richardson DM, Pergl J, Jarošík V, Sixtová Z, Weber E (2008) Geographical and taxonomic biases in invasion ecology. Trends in Ecology and Evolution 23, 237–244.

Pyšek P, Jarošík V, Hulme PE, Pergl J, Hejda M, Schaffner U, Vilà M (2012) A global assessment of invasive plant impacts on resident species, communities and ecosystems: the interaction of impact measures, invading species’ traits and environment. Global Change Biology 18, 1725–1737.

R Core Team (2015) R: A language and environment for statistical computing. R Foundation for Statistical Computing, Vienna, Austria. http://www.Rproject.org.

Russell A E, Raich JW, Vitousek PM (1998) The ecology of the climbing fern *Dicranopteris linearis* on windward Mauna Loa, Hawaii. Journal of Ecology 86, 765–779.

Spotswood EN, Meyer J-Y, Bartolome JW (2012) An invasive tree alters the structure of seed dispersal networks between birds and plants in French Polynesia. Journal of Biogeography 39, 2007–2020.

Steadman DW (2006) ‘Extinction and biogeography of tropical Pacific birds.’ (University of Chicago Press: Chicago, IL, USA)

Stevenson J, Benson A, Athens JS, Kahn J, Kirch PV (2017) Polynesian colonization and landscape changes on Mo’orea, French Polynesia: The Lake Temae pollen record. The Holocene 27, 1963–1975.

Sykes WR (2016) ‘Flora of the Cook Islands.’ (National Tropical Botanical Garden: Kalāheo, HI, USA)

Thaxton JM, Platt WJ (2006) Small-scale fuel variation alters fire intensity and shrub abundance in a pine savanna. Ecology 87, 1331–1337.

Trauernicht, C, Pickett E, Giardina CP, Litton CM, Cordell S and Beavers A (2015) The Contemporary Scale and Context of Wildfire in Hawai’i 1. Pacific Science 69, 427–444.

Venables WN, Ripley BD (2002) ‘Modern Applied Statistics with S.’ Fourth Edition. (Springer: New York, NY)

Vitousek PM, D’Antonio CM, Loope LL, Westbrooks R (1996) Biological invasions as global environmental change. American Scientist 84, 468–478.

Wagner WL, Lorence DH (2002) Flora of the Marquesas Islands website. http://botany.si.edu/pacificislandbiodiversity/marquesasflora/index.htm [Accessed online February 2014]

Washburn, L and Brooks A, of Moorea Coral Reef LTER. 2013. MCR LTER: Coral Reef: Gump Meteorological Data. knb-lter-mcr.9.36 (http://metacat.lternet.edu/knb/metacat/knb-lter-mcr.9.36/lter). Accessed online July 1, 2013.

Whistler AW (2009) ‘Plants of the canoe people: an ethnobotanical voyage through Polynesia.’ (National Tropical Botanical Garden: Kalaheo, HI, USA)

Whistler AW (1996) ‘Wayside plants of the islands: A guide to lowland plants of the Pacific Islands.’ (University of Hawai‘i Press: Honolulu, HI, USA)

